# A multi-view model for relative and absolute microbial abundances

**DOI:** 10.1101/761486

**Authors:** Brian D. Williamson, James P. Hughes, Amy D. Willis

**Author notes:** Corresponding author: Amy D. Willis, Department of Biostatistics, University of Washington, F-600, Health Sciences Building, Box 357232, Seattle, WA 98195-7232.

## Abstract

The absolute abundance of bacterial taxa in human host-associated environments play a critical role in reproductive and gastrointestinal health. However, obtaining the absolute abundance of many bacterial species is typically prohibitively expensive. In contrast, relative abundance data for many species is comparatively cheap and easy to collect (e.g., with universal primers for the 16S rRNA gene). In this paper, we propose a method to jointly model relative abundance data for many taxa and absolute abundance data for a subset of taxa. Our method provides point and interval estimates for the absolute abundance of all taxa. Crucially, our proposal accounts for differences in the efficiency of taxon detection in the relative and absolute abundance data. We show that modeling taxon-specific efficiencies substantially reduces the estimation error for absolute abundance, and controls the coverage of interval estimators. We demonstrate the performance of our proposed method via a simulation study, a sensitivity study where we jackknife the taxa with observed absolute abundances, and a study of women with bacterial vaginosis.

## 1 Introduction

The microorganisms that inhabit a host-associated environment can have a substantial impact on host health (The Human Microbiome Project Consortium 2012, Libertucci & Young 2018, Lloyd-Price et al. 2019). Each microbial taxon present in an environment has a *bacterial concentration*, reflecting the absolute abundance of the taxon per unit volume and therefore the bacterial load on the host. Measuring the concentration of every microbial taxon is resource-intensive, because assays must be designed for each taxon, and it may not be known *a priori* which taxa are present in an environment. It is therefore common to use assays that can detect many taxa; for example, assays based on a hypervariable region of the 16S rRNA gene, or shotgun sequencing of entire microbial communities. While relatively straightforward and inexpensive to perform, these broad range assays can only indicate relative abundances of taxa, and cannot be used to infer absolute abundance (Gloor et al. 2017). However, concentration may be more important than composition for health outcomes, and therefore concentration remains a key quantity of interest in many microbiome studies (Prol et al. 2009, Zemanick et al. 2010, Macé et al. 2013, Stämmler et al. 2016, Vandeputte et al. 2017, Contijoch et al. 2019).

While finding the concentration of every microbe in a highly diverse community is challenging, finding the concentration of a small number of microbes may be tractable. For example, bacterium-specific 16S quantitative PCR (qPCR) assays can be developed on a taxon-by-taxon case (see, e.g., Fredricks et al. 2007, Ryu et al. 2013, Hong et al. 2014). When such data are available, the concentration of a small number of microbes could theoretically be combined with relative abundance data to estimate the concentration of all microbial taxa. A method to accurately estimate all microbial concentrations based on relative abundance data and a small number of microbial concentrations would greatly reduce the labor- and time-intensity of finding the concentration of all microbes in a community. In this paper, we propose and validate a statistical model for this task.

Our approach is to build a hierarchical model that connects the relative abundance data to the absolute abundance data. The observed concentrations of each taxon in each sample are modeled as Poisson-distributed random variables, with taxon- and subject-specific mean parameters that we link to the relative abundances. We observed that 16S sequencing and qPCR assays detected taxa with different efficiencies, and so we incorporate taxon-specific efficiency parameters into our models.

Our paper is structured as follows: the model is defined in Section 2 and estimation is discussed in Section 3. The proposed method is validated on simulated data in Section 4. In Section 5, the proposed estimators are used to estimate the bacterial concentration of all microbes on a study of women with bacterial vaginosis. We provide concluding remarks in Section 6. Software implementing our model and estimators is available in the `R` package `paramedic`, available at github.com/statdivlab/paramedic.

## 2 A model linking absolute and relative abundances

Suppose that we have samples from *n* microbial communities. Let the concentration (absolute abundance in, e.g., gene copies per unit volume or colony-forming units per unit volume) of taxon *j* in community *i* be denoted by *µ*_*ij*_, for *i* = 1,…, *n* and *j* = 1,…, *q*; and 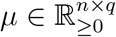 be the matrix of all taxon abundances in all samples. Not all taxa must be present in all communities, and so *µ* may be a sparse matrix.

It is not possible to directly observe *µ* for any taxon because of stochasticity in measuring concentrations (Bonk et al. 2018). However, we are able to obtain realizations from a distribution with expectation *µ*_*ij*_. Unfortunately, sampling from this distribution for all *j* is not typically possible, or is prohibitively expensive. We therefore obtain observed concentrations

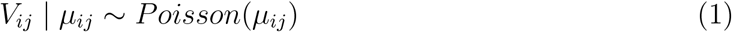

for all *i* but only *j* = 1,…, *q*^obs^, where *q*^obs^ < *q*. It is important to distinguish between the true concentration *µ*_*ij*_ and the observed concentration *V*_*ij*_. Even if *µ* > 0, we may observe a zero concentration in any given sample. Stated differently, a zero observed concentration does not imply that the taxon has zero abundance in the community from which the sample was drawn.

While we are not able to observe concentration data for taxa *j* = *q*^obs^ + 1,…, *q*, we are able to collect relative abundance data for all taxa *j* = 1,…, *q*. Let *W*_*ij*_ be the number of sequencing reads (counts) observed from taxon *j* in sample *i*, and 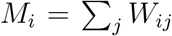 be the total reads observed from sample *i*. A natural model to connect *W*_*i*._:= (*W*_*i*1_,…, *W*_*iq*_) to *µ*_*i*._:= (*µ*_*i*1_,…, *µ*_*iq*_) is

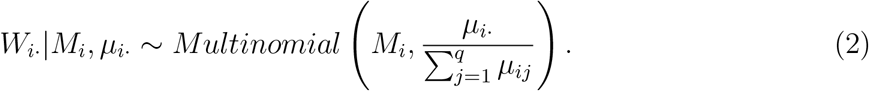

A first-order delta method approximation gives us that under models (1) and (2),

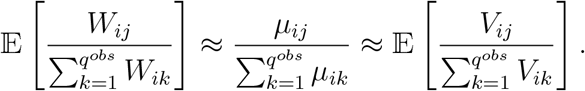

If this approximation holds, we would expect that a scatterplot of 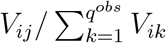 versus 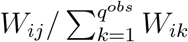 for *i* = 1,…, *n* and *j* = 1, … *q*^*obs*^ would show random scatter around the *x* = *y* line for each taxon. We show this scatterplot in Figure 1 using vaginal microbiome data described in Section 5, and do not observe the expected pattern. Instead, we see that 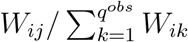 is proportional to 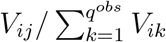, but each taxon has a different slope. This suggests that the model (2) is misspecified in expectation, motivating our proposed model

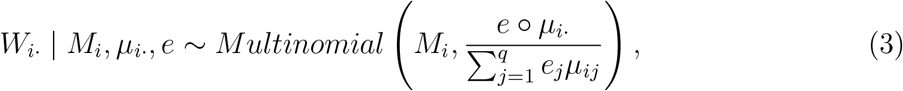

where ◦ denotes Hadamard product (pointwise multiplication), *e*:= (*e*_1_,…, *e*_*q*_), and *e*_*j*_ is the *efficiency* of taxon *j* for being observed by the relative abundance technology compared to the absolute abundance technology. Our efficiency vector *e* plays the role of the “total protocol bias” parameter of McLaren et al. (2019). We now discuss estimation of the parameters of this model, including the identifiability of the efficiencies *e*.

**Figure 1:**
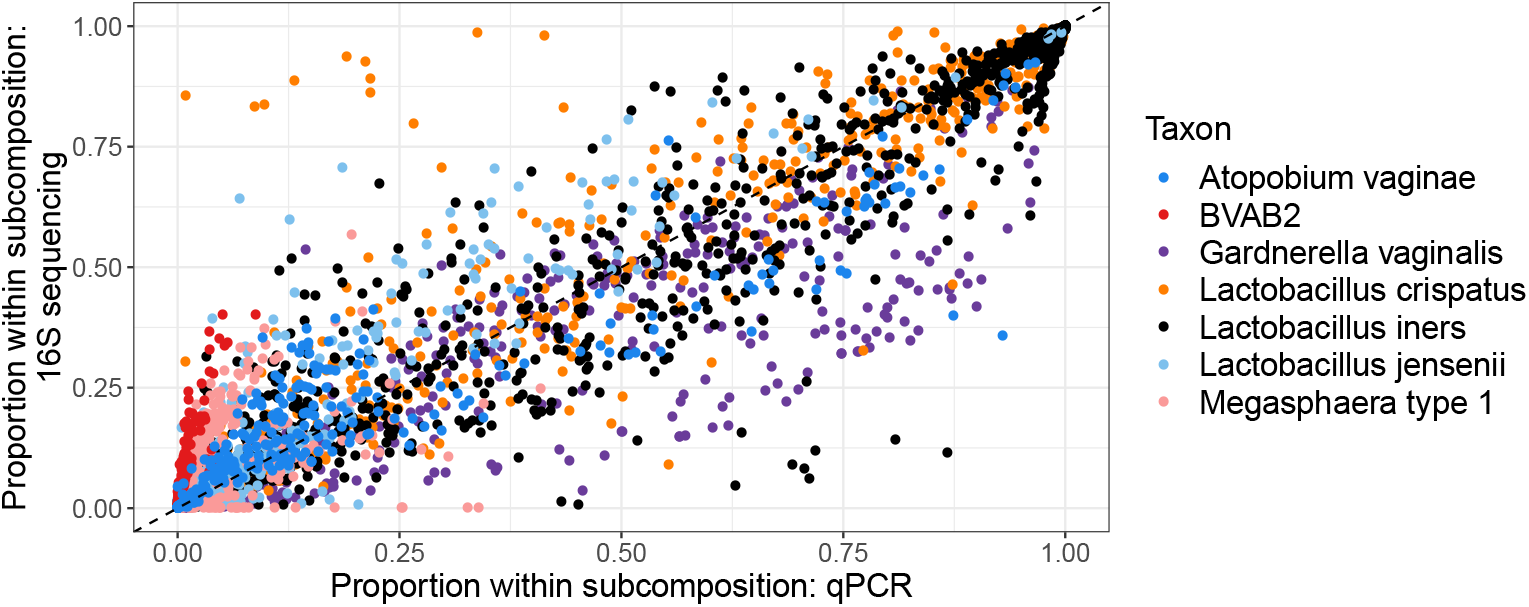
The relative abundance of taxa observed with qPCR versus the relative abundance of the taxa observed by sequencing a hypervariable region of the 16S gene. Note that the subcompositional relative abundance is shown, where the subcomposition is to taxa observed by qPCR. Specifically, 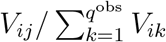 is plotted against 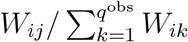. In this dataset, *q*^obs^ = 7 and *n* = 1213. A color version of this figure can be found in the electronic version of the article.

## 3 Estimating model parameters

Our primary goal is to construct point and interval estimators for the *µ*_*ij*_ for all *i* and *j*. A secondary goal is to construct prediction interval estimators for the unobserved concentrations *V*_*ij*_ for all *i* and *j* = *q*^*obs*^ + 1,…, *q*. In this section, we propose three estimation procedures based on the model described in Section 2.

### 3.1 A simple, efficiency-naïve estimator

A simple estimator of *µ*_*ij*_, the concentration of taxon *j* in sample *i*, is

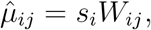

where *s*_*i*_ is a sample-specific scaling factor. This estimate arises naturally because 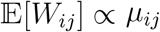. In addition, if *e*_*j*_ for *j* > *q*^obs^ is not estimable, assuming that *e*_*j*_ = *e*_*k*_ for all taxa *j, k* may be necessary. An estimate of the scaling factor could then be obtained by considering the implied scaling factor based on aggregating all observed taxa:

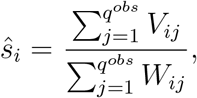

yielding the estimator

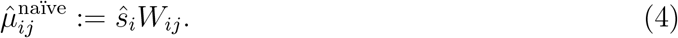

While we did not find a reference to estimator (4) in the literature, it is connected to the proposal of Jian et al. (2018) (see also Vandeputte et al. (2017), Liu et al. (2017), Kevorkian et al. (2018), Gibson & Gerber (2018), Contijoch et al. (2019), Morton et al. (2019)). Jian et al. (2018) consider the problem where the total concentration of all bacteria, 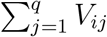, is observed for all *i*, and *W*_*ij*_ is also observed for all *i* and *j*. They wish to estimate *µ*_*ij*_ for all *i* and *j*. Their proposed estimator is 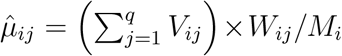. Boshier et al. (2019) recently validated this proposal using taxon-specific qPCR primers and found it to be “predictive of absolute concentration with certain key exceptions,” such as certain taxa and low biomass (low total bacterial concentration: 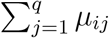) samples. Bonk et al. (2018) give an excellent overview of sources of discrepancies between qPCR and 16S sequencing data.

Previous authors have not proposed methods for quantifying the uncertainty of these naïve estimators. However, interval estimators for 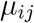 and prediction interval estimators for 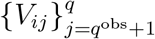 may be constructed by using (1) and (2), the maximum likelihood estimators of the model parameters for *j* ∈ {1,…, *q*^obs^}, and the delta method. To ensure that the interval estimator lies within the parameter space for *µ*, we use a log transformation and estimate

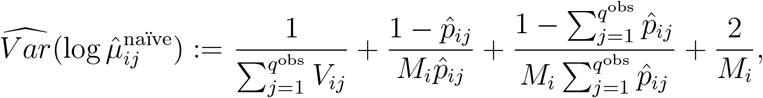

where 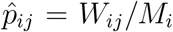. We provide a derivation of this result in the Supporting Information (Section SI 1). A 100(1 − *α*)% confidence interval for *µ*_*ij*_ may then be constructed as

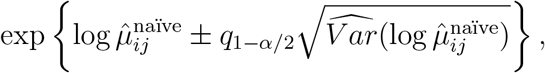

where *q*_*γ*_ is the *γ*-quantile of the standard normal distribution. We can additionally form a 100(1 − *α*)% prediction interval for 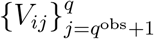:

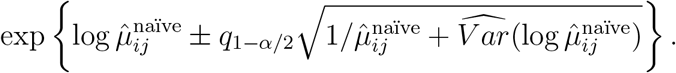

We refer to estimator (4) as the *naïve* estimator because its simplicity must be traded off with its potential drawbacks. First, if the efficiencies are truly unequal, then assuming that the *e*_*j*_’s are equal will lead to biased estimates of *µ*_*ij*_. It will also lead to invalid interval estimates, because the above intervals were constructed under the assumption that the *e*_*j*_’s are equal. Furthermore, these intervals can only be constructed if 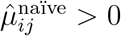, or equivalently, *W*_*ij*_ > 0. However, 16S data is typically very sparse, with *W*_*ij*_ = 0 for many *i* and *j*, and so the naïve interval estimates cannot be constructed for a large fraction of taxa and samples (in our dataset analyzed in Section 5, *W*_*ij*_ = 0 for 95% of the observations). These drawbacks led us to consider more sophisticated estimators, which we now describe. We will benchmark our more flexible proposals against the naïve estimator in our simulation study.

### 3.2 A fully Bayesian estimator with variable efficiency

#### 3.2.1 Parameter point estimation

Bayesian hierarchical modeling is one possible strategy for modeling *V* and *W* to estimate *µ*_*ij*_ and predict *V*_*ij*_ for all *i* and *j*. A hierarchical modeling procedure has several desirable statistical properties here: (i) the joint data model may be specified flexibly; (ii) sampling from the posterior distributions can be performed using freely-available and fast general-purpose software; and (iii) posterior estimates and prediction intervals obtained through this procedure are straightforward to interpret in the context of the generative model. Our goal is to construct valid point and interval estimators in the presence of potentially unequal efficiencies, and when *W*_*ij*_ = 0.

To reflect the differing efficiency with which taxa are detected by 16S and qPCR data (see, e.g., Figure 1) we consider the following model:

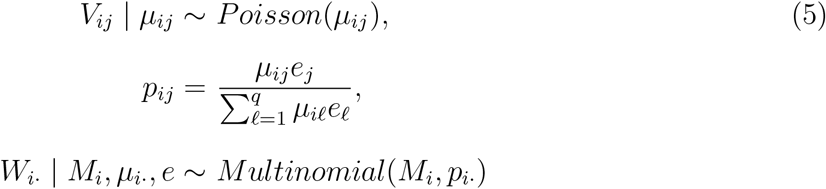

for all *i* and *j*. Next, we must specify the prior distributions of the parameters *µ*_*ij*_ and *e*_*j*_.

Since there is often substantial right skew in the observed *V*_*ij*_ (Section 5), and to ensure positivity of the concentration *µ*_*ij*_, we propose a lognormal prior on the *µ*_*i*._:

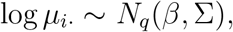

where Σ is a diagonal matrix. We place the following distributions on the hyperparameters *β* and Σ:

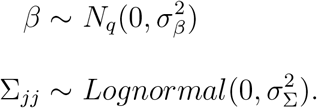

We introduce the following prior on the *e*_*j*_:

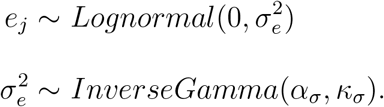

This soft-centering approach makes the parameters *e*_*j*_ and *µ*_*i.*_ identifiable. We note that samples from the posterior distribution of *e*_*j*_ do not satisfy the property that 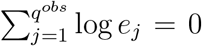 nor that 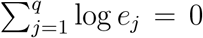 exactly, though we find that both summations are close to zero in practice. We also investigated a hard-centering approach involving the model 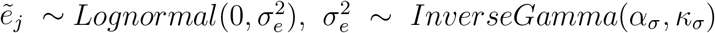, and 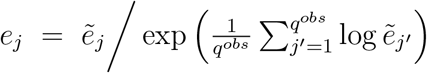. However, we found little difference between the point and interval estimates obtained from the hard- and soft-centering approaches, and similarly for hard-centering over all taxa 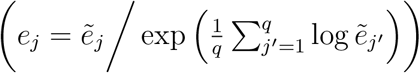. Throughout this manuscript we show results for the soft-centering approach only.

We discuss our default choices of 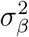, 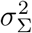, *α*_*σ*_, and *κ*_*σ*_ in Section 4; however, in practice these hyperparameters may be based on independently observed data, numerical experiments, expert opinion, or a combination of these three.

We fit hierarchical model (5) using Stan (Carpenter et al. 2017). Stan is an imperative probabilistic programming language that uses assignment and sampling statements to specify a log-density function. Fully Bayesian inference is available using Hamiltonian Monte Carlo sampling; point estimates may additionally be computed using optimization. Since our parameter space (*µ*, *β*, Σ_11_,…, Σ_*qq*_, 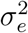) is continuous, Stan is ideal for fitting our model. After fitting the model, we obtain samples from the posterior distribution of (*µ*, *β*, Σ_11_,…, Σ_*qq*_, 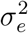).

#### 3.2.2 Interval construction

We now discuss obtaining interval estimates for *µ*_*ij*_ and prediction interval estimates for *V*_*ij*_ using the fitted model. Let 1 − *α* denote the desired level for intervals.

##### Credible intervals for *µ*

100(1 − *α*)% credible intervals for *µ*_*ij*_ are constructed via the (*α*/2, 1 − *α*/2)-quantiles of the posterior sampling distribution of *µ*_*ij*_ based on our proposed hierarchical model.

##### Wald-type prediction intervals for *V*_*ij*_

We incorporate the hierarchical uncertainty of our proposed model into an interval estimate of 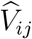 using a Wald-type interval. Using the law of iterated variance conditional on the true *µ*_*ij*_ and our model that *V*_*ij*_ ∼ *Poisson*(*µ*_*ij*_), we estimate the variance in the prediction 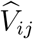 as:

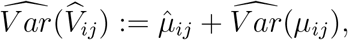

where 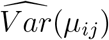 is the variance of the posterior sampling distribution of *µ*_*ij*_ and 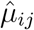 is the posterior mean. Then our prediction intervals for *V*_*ij*_ are

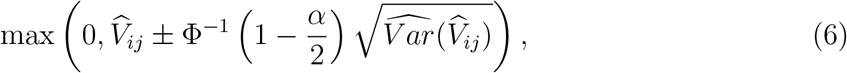

where Φ^*−*1^(*γ*) is the *γ*-quantile of the standard normal distribution. We truncate the lower limit of the prediction interval at zero to reflect that bacterial concentrations are nonnegative.

##### Quantile-based prediction intervals for *V*_*ij*_

We also investigated a quantile-based approach for prediction interval construction, but found its performance to be extremely similar to the Wald-type prediction intervals. We outline the quantile-based approach in Supporting Information (Section SI 2).

#### 3.2.3 An efficiency-naïve estimator

A simplified model may easily be obtained by assuming that all of the efficiencies are equal:

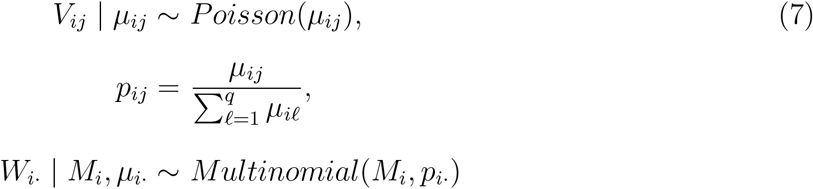

for all *i* and *j*. We use this model in simulated examples to highlight the negative consequences of assuming equal efficiencies when efficiencies are truly unequal. We suggest that model (5) always be used.

#### 3.2.4 Advantages of the varying-efficiency model

Before comparing and validating each of these models and estimators on simulated and observed data, we briefly note some of the advantages of our proposed varying-efficiency model compared to existing and naïve approaches. First, we connect the relative abundance and absolute abundance via a statistical model. Second, by modeling the efficiencies explicitly, we account for the fact that the relative abundances are proportional to the absolute abundances but with a taxon-specific slope, as we observed in Figure 1. Our proposal naturally incorporates the additional uncertainty associated with the unknown efficiencies into our interval estimators. Our relative abundance parameters obey the constraint that 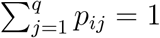 for all *i*. Finally, by adopting a Bayesian hierarchical modeling approach, we can obtain the posterior distribution of *µ*_*ij*_, *j* = *q*^*obs*^ + 1,…, *q*. In other words, we are able to estimate the concentration of taxa for which we do not have absolute abundance data, and construct interval estimators for the concentration of those taxa even when the observed relative abundance is zero. The posterior distribution of the concentration of taxon *j* for *j* > *q*^*obs*^ will be driven by *W*_*ij*_ and *V*_*ij*_ for *j* ≤ *q*^*obs*^, and the prior parameters 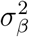, 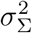, *α*_*σ*_, and *κ*_*σ*_. We note that the interval estimates for *µ*_*ij*_ and *V*_*ij*_ can be wide for *j* > *q*^*obs*^.

## 4 Results under simulation

We now present simulation results on the performance of the estimators proposed in Section 3. In all cases, we use Stan to fit hierarchical models (5) and (7) using 4 chains per simulated dataset, each with 10,000 burn-in iterations and 10,500 total iterations (2,000 total iterations for each of *B* = 50 simulations for each set of parameters to investigate). We describe our process for initializing these chains in the Supporting Information (Section SI 3). It was not feasible to confirm convergence for every individual simulation via trace plots, and so we confirmed that the median and interquartile range (IQR) of the Gelman-Rubin 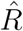 statistic (Gelman & Rubin 1992) was close to 1 for all parameters of interest. All code for replicating the simulation results is available at github.com/bdwilliamson/paramedic_supplementary.

For each of *b* = 1,…, *B* Monte Carlo samples (repeated simulations), we assess performance using:

i. Root mean squared error for *µ*_*ij*_, averaged across all *n* samples and *q* taxa:

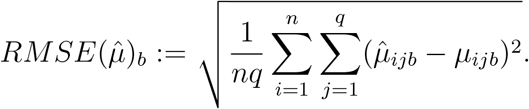
ii. Root mean squared prediction error (RMSPE) for *V*_*ij*_, *j* = *q*^*obs*^ + 1,…, *q*, averaged across all *n* samples and *q* − *q*^obs^ unobserved taxa:

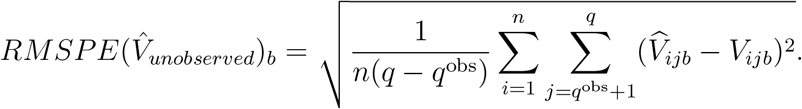
iii. Average coverage of 95% posterior credible intervals for *µ*_*ij*_: Let 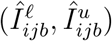 be the proposed credible interval for *µ*_*ij*_ in the *b*th simulation. Then

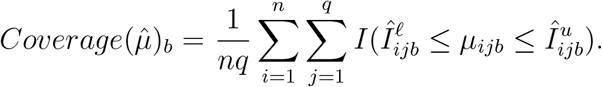
iv. Average coverage of 95% posterior prediction intervals for *V*_*ij*_, *j* = *q*^*obs*^ + 1,…, *q*: Let 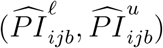 be the proposed posterior prediction interval for *V*_*ij*_ in the *b*th simulation. Then

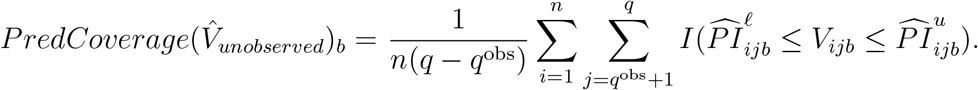

While our primary goal is estimation of the true concentration *µ*, we also investigate the performance of predicting *V* for the unobserved taxa, as this may be of interest in some settings (e.g., for assessing correct model specification; see Section 5.3 and Figure 5).

We report these four summaries for each estimator under consideration. In each case, we display the average of the summary measure over Monte Carlo replicates. In all simulations, we exclude taxa whose mean expected abundance *µ*_*ij*_, averaged over all samples, is below 1 unit. In practice, taxa observed in low abundance across all samples are typically excluded from analysis (Callahan et al. 2016), and so this reflects the typical use case of the proposed method. However, in practice *µ*_*ij*_ is unknown, and thus exclusion may be done based on *W*_*ij*_. We provide a discussion of filtering rules and the rationale behind the particular rule used here in the Supporting Information (Section SI 4.2). Finally, if the naïve estimate for a given sample and taxon is zero, then we do not include that sample-taxon pair when computing average coverage of naïve interval estimates.

### Default parameters

We strongly recommend that the user investigate the sensitivity of results to prior parameters. Additionally, the values of prior parameters should be carefully chosen to match the measurement scale of the dataset. In our dataset of Section 5, the sample variances of the realized log-qPCR data are near 50. Based on this observation, we chose 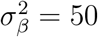 and 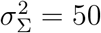 as default parameters for our simulation study. We additionally chose *α*_*σ*_ = 2 and *κ*_*σ*_ = 1 since these choices led to fast convergence of our sampling algorithm in our data analysis. We provide an investigation of the prior parameters 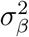 and 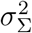 in the Supporting Information (Section SI 4.3).

### Simulation settings

Throughout this section we simulate data according to *M*_*i*_ ∼ DiscreteUniform(10^4^, 10^5^), reflecting the distribution of read depths that we observed in our data. We also simulate data according to 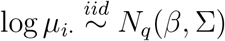 for all subjects *i* = 1,…, *n* where 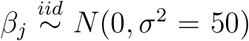 for all *j* and Σ = **I**_**q**_. In all cases, we simulate *V*_*ij*_ ∼ *Poisson*(*µ*_*ij*_) and *W*_*i.*_ ∼ *Multinomial*(*M*_*i*_, *p*_*i.*_), where 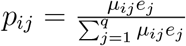. The specific choices for the distribution of *e*_*j*_ and the values of *q* and *q*^*obs*^ vary in each simulation.

#### 4.1 Effect of varying the number of taxa

We first investigate the effect of varying *q* and *q*^obs^ while holding other parameters fixed. We simulated data with no varying efficiency (*e*_*j*_ = 1 for all *j*) and fit the efficiency-naïve model (7). We investigate the efficiency model in Section 4.2.

We observe 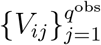 and 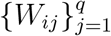 for *i* = 1,…, *n*, where *n* = 100. We vary *q* ∈ {10, 20, 40, 60}; for each *q*, we additionally vary *q*^obs^ ∈ {2, 3,…, 7}. For each unique combination of *q* and *q*^obs^, we generate data from this population by: (i) generating *β* and Σ; and (ii) generating *µ*_*ij*_, *V*_*ij*_, *M*_*i*_, and *W*_*ij*_. We then repeat step (ii) to obtain *B* = 50 independent Monte-Carlo replicates.

In microbiome studies, qPCR data is typically available for only the taxa that are of most interest to the investigator or are expected to be most abundant. For this reason, in our simulations the *q*^obs^ most abundant taxa based on the observed *W*_*ij*_, averaged over the *n* samples, are used to estimate *µ* for all taxa and predict the unobserved qPCR data, 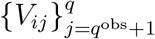. This means that in our simulations, as *q* increases we add increasingly rare taxa.

Figure 2 displays the results of this experiment. In the top row, we see that nominal 95% credible intervals for *µ* based on the naïve estimator have slightly greater average coverage than our proposed estimator. However, the average coverage of the proposed credible intervals for *µ* is close to nominal for all (*q, q*^obs^) combinations. We note that for both estimators, average coverage for *µ* decreases as *q* increases for a fixed *q*^obs^, but increases as *q*^obs^ increases for a fixed *q*. The decrease in average coverage as *q* increases is due to poor marginal coverage for the lowest abundance taxa (see Supporting Information, Section SI 4.4). We also see that average coverage of prediction intervals for *V* based on the proposed estimator is at the nominal level for all (*q, q*^obs^) combinations. This is encouraging, especially in view of the fact that we often have many more relative abundance measurements than species-specific qPCR measurements; indeed, the results we present in Section 5 are based on *q*^obs^ = 7. In contrast, average coverage of prediction intervals based on the naïve estimator is below the nominal level for large *q*. This poor performance of intervals based on the naïve estimator is due in large part to the fact that a naïve interval does not exist when the naïve estimator equals zero. The proportion of cases where the naïve estimator is zero, and thus excluded from computing performance, is 0.17%, 1.5%, 26%, and 50% of sample-taxon pairs for *q* = 10, 20, 40 and 60, respectively. Additionally, since we compute intervals based on the naïve estimator on the log scale, the lower limit of the back-transformed interval is almost surely greater than zero, if the interval exists. This leads to under-coverage of cases where the true qPCR value is exactly zero, which is increasingly the case as *q* increases.

**Figure 2:**
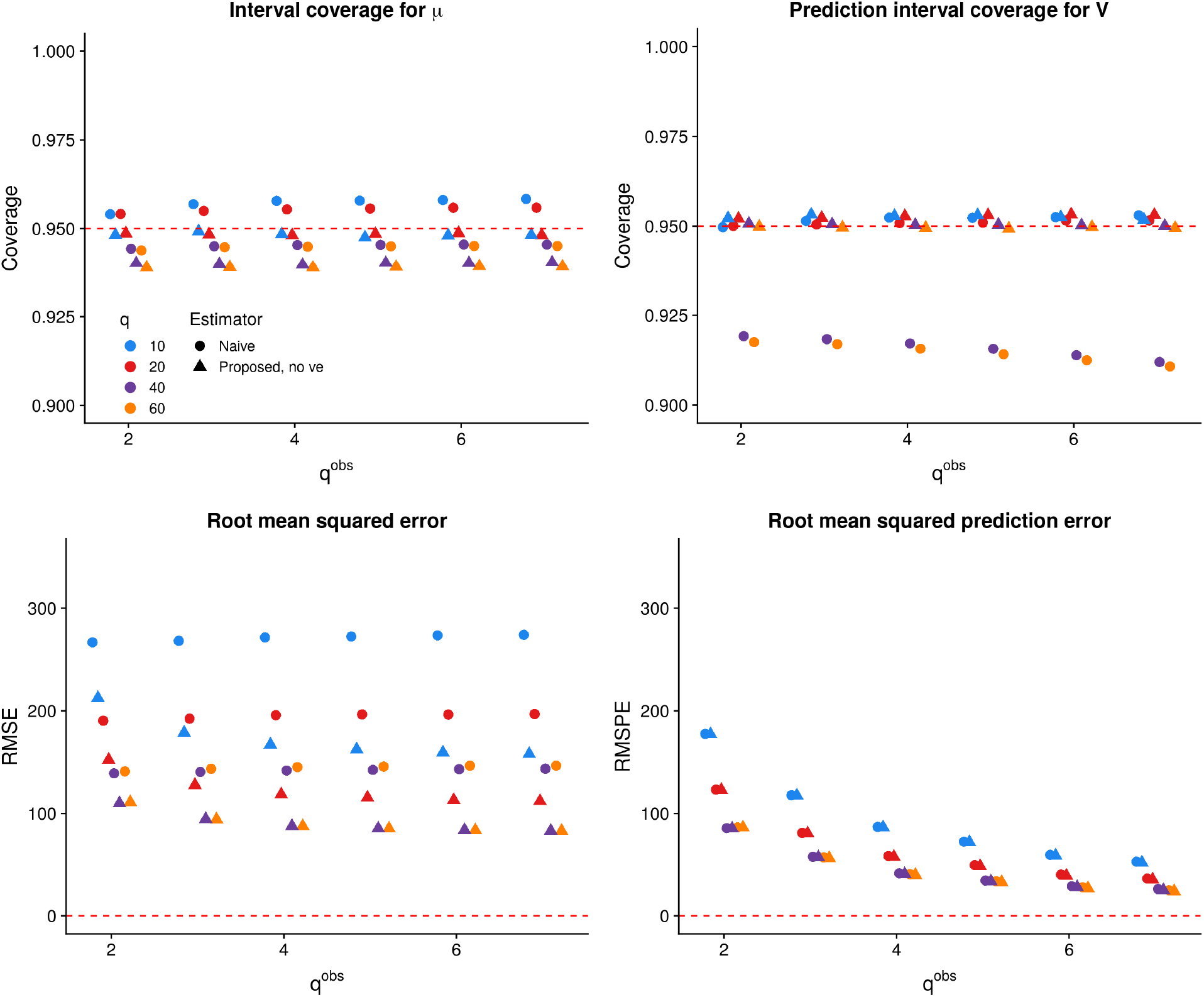
Performance of the naïve estimator and proposed estimator with no varying efficiency versus *q*^obs^ for *q* ∈ {10, 20, 40, 60}. *Top row*: coverage of nominal 95% intervals based on the naïve (circles) and proposed (triangles) estimators. *Bottom row*: root mean squared prediction error and root mean squared error for both estimators. In each plot, color denotes *q*, while shape denotes the estimator. A color version of this figure can be found in the electronic version of the article.

In the bottom row of Figure 2, we see that the proposed estimator has lower RMSE than the naïve estimator over all (*q, q*^obs^) combinations, while the RMSPE of the two estimators is comparable. As *q*^obs^ increases for a fixed *q*, both RMSE and RMSPE tend to decrease. We provide evidence in Section SI 4.1 that the proposed estimator has low bias, and thus the RMSE of the proposed estimator appears to be driven by its variance.

After averaging over Monte-Carlo replicates, the median Gelman-Rubin 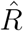 for *µ* over all samples and taxa for *q* = 60 and *q*^obs^ = 7 was 0.99, with an inter-quartile range of [0.99, 1.00], showing excellent convergence; convergence was similar in other pairings of *q* and *q*^obs^ and for *β* and Σ for each pairing. We investigated the trace plots for a small number of Monte Carlo samples, which showed well-mixed chains after the burn-in period.

In many experiments, *q* may be much larger than 60. For example, in our data analysis of Section 5, *q* = 433. We anticipate that the trends observed in this simulated experiment would hold for larger *q*, but did not investigate them here because the time required to compute our estimator increases with *q*.

#### 4.2 Varying the distribution of efficiency

In this experiment, we fix *q* = 40 and *q*^obs^ = 7. We vary *σ*_*e*_ ∈ {0, 0.1,…, 0.5, 0.6, 0.8, 1}. For each *σ*_*e*_, we generate data from this population in the same manner as the previous experiment, resulting in 50 independent Monte-Carlo replicates. We use Stan to fit our proposed variable efficiency model (5) and our efficiency-naïve model (7). We refer to the proposed estimator that does not model varying efficiency as “proposed, no ve”, while we refer to the proposed estimator that does model varying efficiency as “proposed, ve”. As we have described before, the naïve estimator does not account for varying efficiency.

Figure 3 displays the results of this experiment. In the top row, we see that as *σ*_*e*_ increases, the prediction interval average coverage and credible interval average coverage decline to levels below 95% for the naive and efficiency-naïve Bayesian models but are maintained close to or above 95% for the proposed varying efficiency model. This coincides with our expectation that varying efficiency must be modeled if it is truly present. In the bottom row, we see that as *σ*_*e*_ increases, the RMSE and RMSPE of all three estimators increases. The proposed estimator that does model varying efficiency tends to have the lowest RMSE and the highest RMSPE; however, as *σ*_*e*_ increases, the difference between the RMSPE of the estimators decreases. Since we observed nearly identical patterns for the same experiment with *q*^obs^ = 3, we do not show those results here. In the data we analyze in Section 5, we estimate 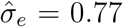. This suggests that interval estimates based on the proposed estimator will be more reliable with respect to interval coverage on this dataset.

**Figure 3:**
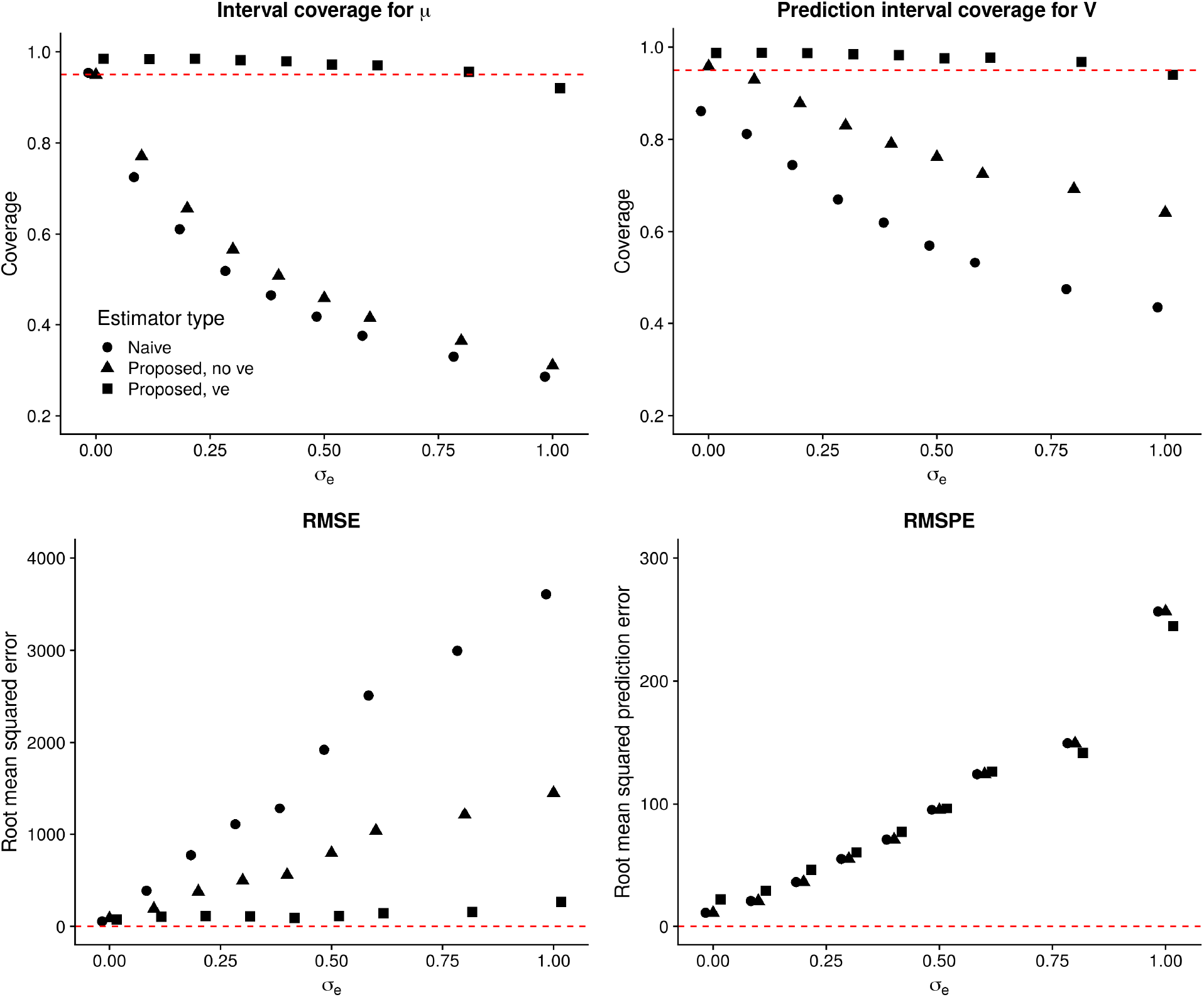
Performance of the naïve estimator and proposed estimator without and with varying efficiency versus *σ*_*e*_ for *q* = 40 and *q*^obs^ = 7. *Top row*: coverage of nominal 95% intervals based on the naïve (circles) and proposed estimators without (triangles) and with (squares) varying efficiency. *Bottom row*: root mean squared prediction error and root mean squared error for both estimators.

After averaging over Monte-Carlo replicates, the median Gelman-Rubin 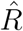 for *µ* over all samples and taxa for *σ*_*e*_ = 0.5 was 1.00 (IQR [0.99, 1.00]) when varying efficiency was modeled and 0.99 (IQR [0.99, 1.00]) when efficiency was not modeled. As we varied *σ*_*e*_, the median 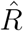 for all model parameters tended to be near one, with a maximum of 1.2 for *β* when *σ*_*e*_ = 0 and varying efficiency was not modeled. Inspection of trace plots for a small number of samples showed well-mixed chains after the burn-in period.

## 5 Results from a study of the vaginal microbiome

### 5.1 Description of the study sample

These data are from Project 2 of the Sexually Transmitted Infections Cooperative Research Center at the University of Washington (UW STICRC). Twenty women were recruited by the UW STICRC because they had a history of frequent episodes of bacterial vaginosis over the past year. Each participant self-collected a vaginal swab every eight hours for 60 days. However, not all participants completed the full sampling schedule, resulting in 1328 observations with 16S relative abundance data. These data are additionally described by Boshier et al. (2019). We use these data for illustrative purposes only, and did not incorporate correlation between samples into our analysis.

The data contain observed concentrations from qPCR (measured in 16S gene copies per swab) on *q*^*obs*^ = 7 taxa, and observed 16S relative abundance data on *q* = 433 taxa. The taxa with observed qPCR data are *G. vaginalis*, *L. crispatus*, *L. iners*, *L. jensenii*, *Megasphaera type 1*, *BVAB2 spp.*, and *A. vaginae*. We removed any sample *i* where *M*_*i*_ was less than 1,000 (outlying low sequencing depth), resulting in a final analysis dataset with *n* = 1213 samples. The overall goals of this analysis were to: (i) estimate the true concentrations *µ* for a subset of the taxa and each sample, and compute valid interval estimates for the true concentrations; and (ii) to predict the bacterial concentrations for taxa *j* > *q*^*obs*^ in each sample, and compute valid prediction interval estimates. We restrict our attention to a subset of the 433 total taxa to reflect limits on computation time and computing memory (see Section SI 5 for details on compute requirements). In all cases, we use Stan to fit our models using four chains, each with 14,000 burn-in iterations and 16,000 total iterations. We also use the default parameters discussed in Section 4.

### 5.2 Results of the primary analysis

In our primary analysis, we analyzed all 1213 samples. Our goal is to estimate concentrations for *q* = 17 taxa.

Figure 4 displays the results of our primary analysis. Panel A (left) is a heatmap showing the estimated log concentrations for the 17 taxa and a random subset of 40 samples, normalized against the maximum estimated concentration in this subsample of samples and taxa. Red denotes large normalized log concentration, while blue denotes small normalized log concentration. We observe substantial variability in concentrations both between samples and between taxa. For example, while *L. iners* appears to be a high-abundance taxon on average, some samples (e.g., samples 1 and 40) have much smaller normalized concentration. This pattern appears more striking in the taxa lacking qPCR measurements: some samples have a large estimated abundance of *BVAB1 spp*. (e.g., subjects 5 and 14), while many others have a low estimated abundance of this same taxon.

**Figure 4:**
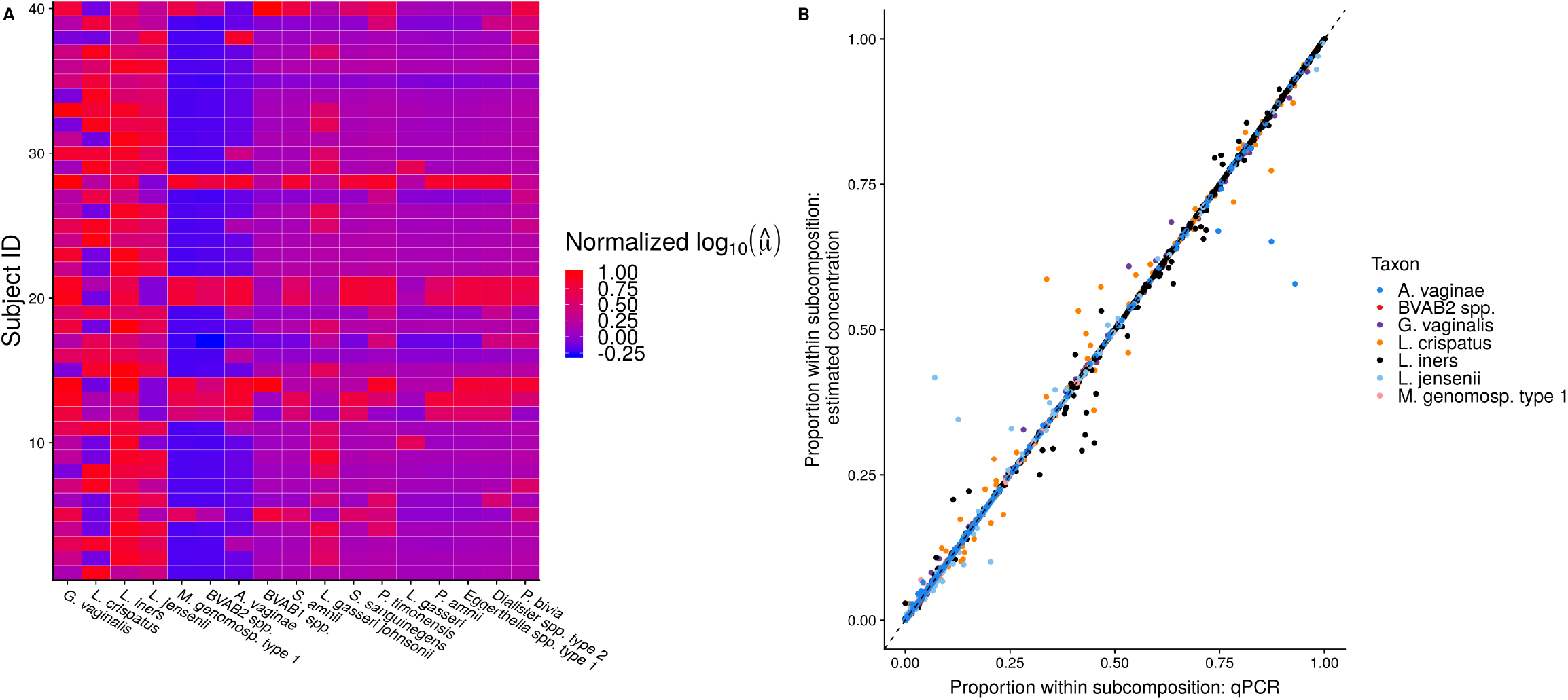
**A**. A heatmap showing normalized estimated log concentrations for the 17 taxa and 40 randomly sampled samples. Red indicates large concentration relative to the maximum in this subsample, while blue indicates small concentration relative to the maximum in this subsample. **B**. The relative abundance of taxa observed with qPCR versus the estimated relative abundance of the taxa based on the variable-efficiency estimator. Specifically, 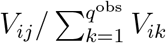 is plotted against 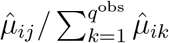. *q*^obs^ = 7 and *n* = 1213 in this dataset. A color version of this figure can be found in the electronic version of the article.

Panel B (Figure 4, right) plots 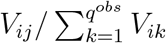 against 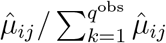. Here, we see that the model accounting for varying efficiency results in good estimates of *µ* on the individual taxa for which we have qPCR data. In the next section, we investigate performance where we do not have qPCR data in a leave-one-out analysis. Finally, we estimate that 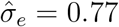, with a 95% credible interval of (0.31, 1.81). We additionally estimate a range of efficiencies among the taxa with observed qPCR of between 0.36 and 6.26. These results together imply that there is substantial variation in taxon efficiencies, and that modeling this variation is important.

We provide additional results in a comma-separated file in Supplementary Data, consisting of: point estimates and credible intervals for *µ*_*ij*_, *i* = 1,…, 1213 and *j* = 1,…, 17; and prediction intervals for *V*_*ij*_, *i* = 1,…, 1213 and *j* = 8,…, 17.

### 5.3 Leave-one-out analysis to predict observed qPCR

To validate our proposed method on these data, we performed a leave-one-out analysis only on those taxa with observed concentrations from qPCR. In this analysis, we first restricted the dataset to only those taxa with observed concentrations, leaving us with 7 taxa with both concentration and relative abundance data. We next randomly selected *n* = 100 samples to limit computation time. Then, for each *k* ∈ {1, 2,…, 7} denoting the taxon to predict, we removed taxon *k* from the observed qPCR matrix, and computed the three estimators of *µ*_*ik*_ (naïve; proposed, no ve; and proposed, ve) and predictions for *V*_*ik*_, as well as prediction intervals for *V*_*ik*_. We then calculated mean squared prediction error and average coverage of prediction intervals (averaging over *i* = 1,…, 100), comparing the estimates of concentration to the observed qPCR concentration.

Figure 5 displays the prediction interval coverage and MSPE for the left-out taxon. Prediction interval coverage of the proposed varying-efficiency estimator is close to or higher than nominal for Wald-type intervals for 6 out of 7 left-out taxa. Furthermore, for 4 out of 7 left-out taxa, the RMSPE is comparable across the three proposed estimators. When *L. crispatus* is left out, the variable efficiency model has lower RMSPE than the nonvariable efficiency models, while when *L. iners* and *A. vaginae* are left out, the nonvariable efficiency models have lower RMSPE. However, the proposed variable efficiency model controls coverage for 6 out of 7 left-out taxa, while the nonvariable efficiency models control coverage in 0 out of 7 cases. This reinforces the importance of modeling varying efficiency: while we saw in Figure 4 that the varying-efficiency model provides valid estimators for the taxa for which we have qPCR data, the model only provides valid estimators on average over all taxa.

**Figure 5:**
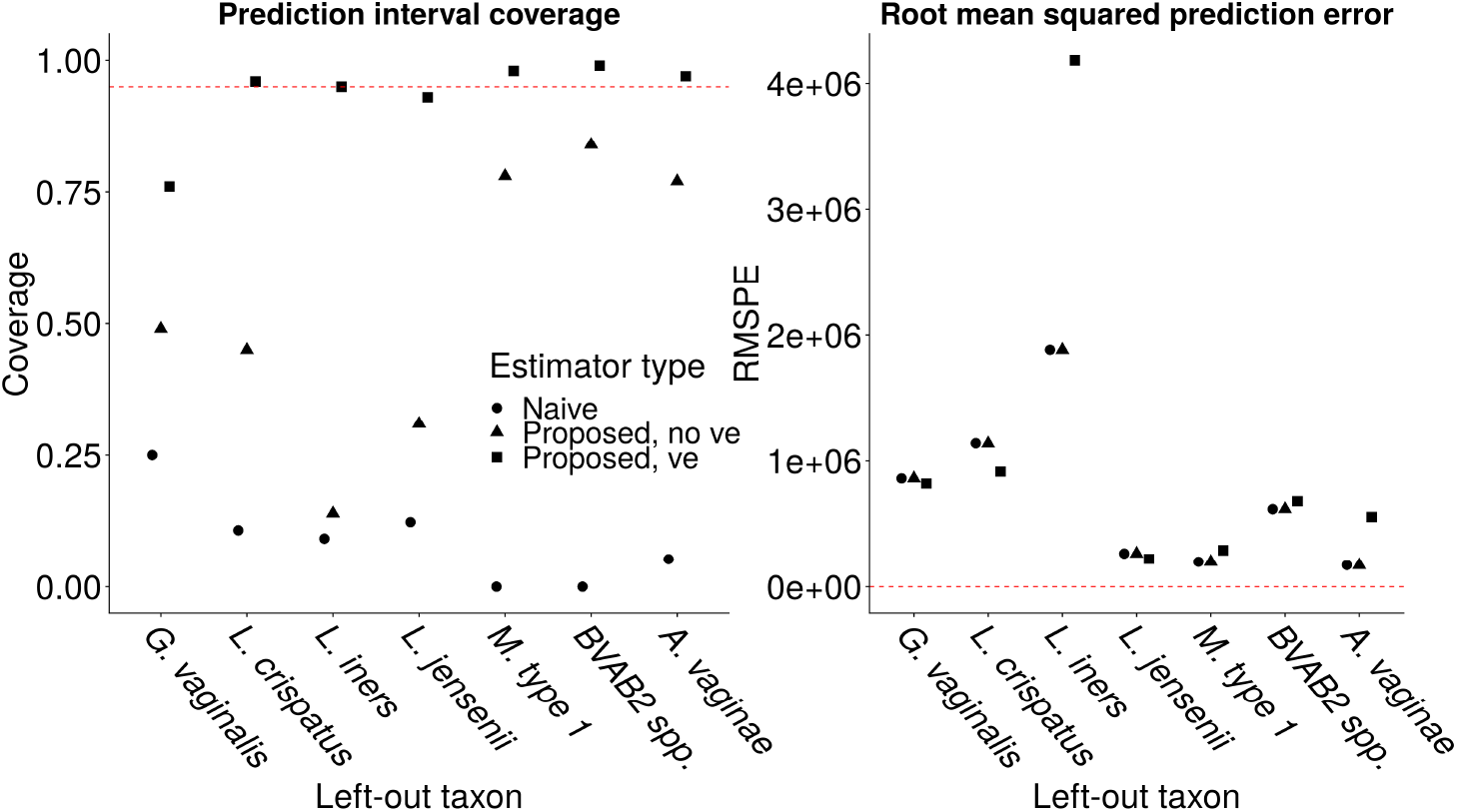
*Left:* Average coverage of nominal 95% prediction intervals for the left-out taxon averaged over study participants. *Right:* MSPE on the left-out taxon. Circles denote the naïve estimator, triangles denote the Bayesian efficiency-naïve estimator, and squares denote the proposed estimator that does model varying efficiency

We conclude by investigating the robustness of the estimators of efficiency to the inclusion of additional qPCR data. In Figure 6, we contrast the distribution of the estimated efficiencies in an analysis with all seven taxa (the full-data analysis) against an analysis with a taxon left out. In the left-hand panel, we leave out *G. vaginalis*; in the right-hand panel, we leave out *BVAB2 spp.* We see in the left-hand panel that the distributions of efficiency for all taxa are nearly identical between the leave-one-out analysis and the full-data analysis, except that the distribution of *G. vaginalis* regresses to the mean and increases in variance when that taxon is left out. This indicates that *G. vaginalis* is a low-efficiency taxon. Note that the median estimated efficiency is close to the prior mean value in the leave-one-out analysis. We see the same pattern of regression to the mean and increase in uncertainty when *BVAB2 spp.* is left out. The inclusion of *BVAB2 spp.*, which is a high-efficiency taxon, alters the estimated efficiencies of the remaining taxa, resulting in increased estimated variance. These results indicate that the algorithm learns differently based on which taxa are observed: if a high efficiency taxon (e.g., *BVAB2 spp.*) is observed in both the absolute and relative abundance data, then the algorithm detects this larger variance in the efficiencies. This reinforces that even a model designed to account for the distribution of varying efficiencies cannot accurately predict the efficiency of an individual taxon when only relative abundance data are available. Note that these findings corroborate existing literature: Boshier et al. (2019) found that *BVAB2 spp.* is a high-efficiency taxon, and McLaren et al. (2019) found that *G. vaginalis* is a low-efficiency taxon.

**Figure 6:**
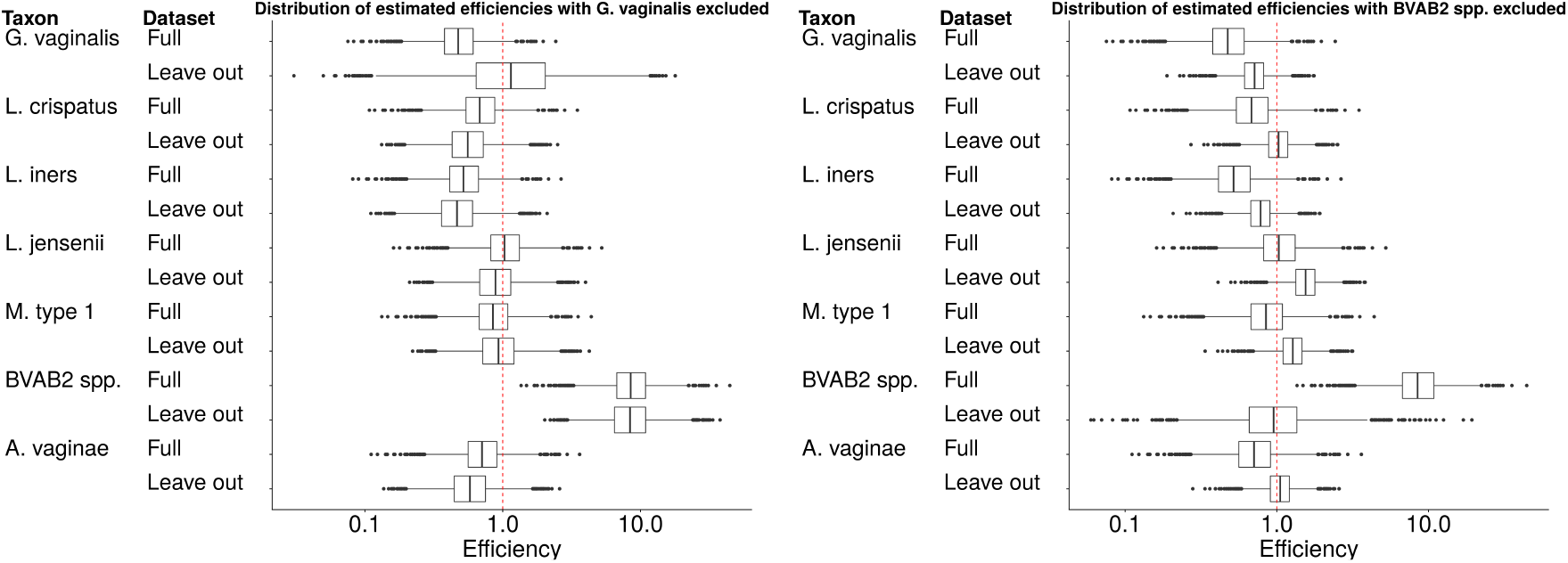
Boxplots showing the posterior distribution of estimated efficiencies. *Left:* estimated efficiencies from the full data analysis and from an analysis where *G. vaginalis* was left out. *Right:* estimated efficiencies from the full data analysis and from an analysis where *BVAB2 spp.* was left out.

## 6 Discussion

In this manuscript we developed a statistical procedure for jointly modeling absolute and relative abundance data, with a specific application to qPCR and 16S data collected on microbial communities. We proposed a hierarchical model with the following appealing characteristics: (i) point and interval estimators for the true and realized absolute abundances can be constructed for all taxa and all samples; (ii) average coverage of credible and prediction intervals is controlled at or above the nominal level; (iii) the efficiency of taxon detection of the different technologies is explicitly modeled and allowed to vary; and (iv) the method is implemented as an open-source R package. We observed that estimators based on the proposed model have lower MSE compared to more naïve estimators, and approximately equal MSPE. To our knowledge, our proposed hierarchical model is the first statistical model for this microbial multi-view data structure.

We found strong evidence for differing efficiency of taxon detection between qPCR and 16S technologies. Given that the collection of qPCR data involves calibration (via a “standard curve”) and 16S relative abundance data does not usually involve any calibration, we modeled the efficiency of the 16S data compared to the qPCR data, rather than the efficiency of the qPCR data compared to the 16S data. This is consistent with recent literature highlighting the lack of calibration and resulting bias of 16S data for estimating relative abundance (McLaren et al. 2019). Our method can also be used with other technologies for obtaining absolute and relative abundance data. For example, data from plate counting or flow cytometry could replace qPCR data, and a different taxonomically informative marker could replace 16S sequencing. We note that the default parameters in our software were chosen based on the units and scale of the UW STICRC data, and should be adjusted to reflect the data under study. We found that interval estimates for concentration can be very wide when qPCR primers are unavailable (Figure 6).

Empirically, we found that modeling the efficiency of the different technologies is critical for obtaining accurate estimates of microbial abundance. Boshier et al. (2019) found that a naïve approach consistently overestimates the concentration of certain taxa by an order of magnitude (e.g. *BVAB2*). By using a leave-one-out approach, we observed that modeling varying efficiency achieves near-nominal coverage of prediction intervals, while failing to model varying efficiency does not control coverage (Figure 5). Variation in efficiency between taxa implies that while our method controls coverage on average across all taxa, these properties are not guaranteed for each individual taxon. Incorporating uncertainty in the efficiencies into interval estimators results in wider intervals for the true microbial concentration, but because coverage is controlled, it accurately reflects the level of uncertainty in estimating absolute abundance. We believe that modeling efficiency is a significant advantage of our method over other proposals in the literature for combining relative and absolute abundance data (Liu et al. 2017, Vandeputte et al. 2017, Jian et al. 2018, Contijoch et al. 2019).

Our method involves estimating *n* × *q* concentration parameters *µ*_*ij*_, and *q* efficiency parameters *e*_*j*_. The inclusion of additional samples therefore increases the number of parameters to estimate (a Neyman-Scott problem (Neyman & Scott 1948)). Additionally, for small *q*^obs^ the prior distribution on the efficiencies will play a large role in determining the width of interval estimates for the concentrations *µ*_*ij*_. For these two reasons, instead of increasing *n* or *q*, *q*^*obs*^ should be increased where possible to reduce interval width (see Figure 2). Varying the prior parameters *α*_*σ*_ and *κ*_*σ*_ also alters the width of intervals. Future modeling work could link the concentrations *µ*_*ij*_ to covariates *X*_*i*_ ∈ ℝ^*p*^ to reduce the number of parameters in the model, to model the correlation structure between taxa (see Gibson & Gerber (2018)), and to use additional data on the total bacterial load, 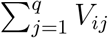, to improve estimates of *µ* and *V* using our proposed varying efficiency model. Future data analysis could compare the estimated efficiencies obtained using different sample preparation and sequencing protocols to select between different experimental protocols.

The proposed method provides a general approach for jointly modeling absolute and relative abundance data where each taxon’s detection efficiency differs across the two data sources. Note that our approach to modeling efficiency can model any multiplicative scaling factor between the data sources, including gene copy number. However, our motivating data sources were 16S community profiling, and taxon-specific qPCR targeting the 16S gene. Because both methods targeted the same gene, our efficiency estimators are not estimating 16S copy number. In the case that different amplicons were targeted and copy numbers were known, copy number differences could be explicitly included with a minor modification to our proposed procedure.

The methods proposed in this manuscript are all implemented in the R package paramedic (<u>P</u>redicting <u>A</u>bsolute and <u>R</u>elative <u>A</u>bundance by <u>M</u>odeling <u>E</u>fficiency to <u>D</u>erive <u>I</u>ntervals and <u>C</u>oncentrations) available at github.com/statdivlab/paramedic. All experiments and data analyses may be replicated using code available at github.com/bdwilliamson/paramedic_supplementary.

## Acknowledgments

The authors thank Ben Callahan, David Clausen, Michael McLaren and Joshua Schiffer for many insightful comments and suggestions that greatly improved the manuscript. The authors also thank David Fredricks for generating the UW STICRC data; see Boshier et al. (2019) for details.

This work was supported by the National Institute of Allergy and Infectious Diseases (NIAID) of the National Institutes of Health (NIH) under award number U19 AI113173, and the National Institute of General Medical Sciences (NIGMS) of the NIH under award number R35 GM133420. The opinions expressed in this article are those of the authors and do not necessarily represent the official views of the NIAID, the NIGMS or the NIH.

## Notes

https://github.com/statdivlab/paramedic/

